# The genomic landscape of early stage ovarian high grade serous carcinoma

**DOI:** 10.1101/2021.05.05.440631

**Authors:** Z. Cheng, H. Mirza, D.P. Ennis, P. Smith, L. Morrill Gavarró, C. Sokota, G. Giannone, T. Goranova, T. Bradley, A. Piskorz, M. Lockley, the BriTROC-1 Investigators^, B. Kaur, N. Singh, L.A. Tookman, J. Krell, J. McDermott, G. Macintyre, F. Markowetz, J.D. Brenton, I.A. McNeish

**Author notes:** to whom correspondence should be addressed: Professor Iain McNeish, Imperial College London, IRDB Building, Hammersmith Hospital, W12 0NN. equal contribution (H. Mirza, D.P. Ennis). The BriTROC Investigators are listed in the Supplementary information. **Statement of translational relevance** To determine whether early stage ovarian high grade serous carcinoma (HGSC) represents a distinct sub-population, we collected samples from 45 patients with early stage HGSC to identify potential short genomic variants and copy number differences, with comparison to a cohort of 51 late-stage cases. We found no significant differences in somatic mutations or focal copy number (CN) between early stage and late stage cohorts. There was, however, a significant difference in both ploidy and copy number signature exposures between early and late stage samples, with greater ploidy and signature 4 exposure in late stage cases. Unsupervised hierarchical clustering revealed three clusters that were prognostic. Together, our data suggest that early and late stage HGSC share fundamental genomic features but that late stage disease appears to evolve from early stage, with the appearance of high ploidy and whole genome duplication.

## Abstract

**Purpose:** High grade serous carcinoma (HGSC) is the commonest type of ovarian cancer. Nearly all HGSC cases are diagnosed at late stage and it is not clear whether early stage HGSC has unique characteristics compared to late stage tumours.

**Experimental Design:** We analysed samples from 45 patients with FIGO stage I - IIA HGSC - 40 from the pathology archives of three large UK cancer centres and 5 from the BriTROC-1 study. We performed shallow whole genome sequencing (sWGS) and targeted next generation sequencing to investigate somatic mutations and copy number alterations. We compared results to 51 stage IIIC/IV HGSC patients from the BriTROC-1 study.

**Results:** There was no difference in median age between the early stage (median 61.3 years, range 40-84) and late stage (median 62.3 years, range 34-76) patients at diagnosis. *TP53* mutations were near-universal (92% early stage, 100% late stage samples) and there were no significant differences in the rates of other somatic mutations, including *BRCA1* and *BRCA2*, or focal copy number alterations between early- and late-stage cohorts. There were also no unique amplifications or deletions in either cohort. However, median ploidy was greater in late stage (median 3.1) than early stage (median 2.0) samples. In addition, there were higher numbers of breakpoints per 10MB and per chromosome arm and higher absolute copy number in late stage than early stage cohorts; early stage samples had longer segment length. Overall copy number signature exposures were significantly different between early and late stage samples with greater signature 3 exposure in early stage and greater signature 4 in late stage. Both simplex plot and unsupervised hierarchical clustering suggested that a subset of late stage samples retain early stage appearances with high signature 3 and co-clustering with the early stage samples

**Conclusions:** These data suggest that there are no unique mutations or focal copy number alterations in early stage HGSC. However, whole genome duplication is significantly more common in late-stage disease, suggesting evolution during disease progression. However, a subset of late stage HGSC retains early-stage features, which are associated with improved overall survival.

## Introduction

High grade serous carcinoma (HGSC) accounts for approximately 70% of all ovarian cancer (OC) cases and approximately 80% of OC deaths. The large majority of patients with HGSC present with advanced (FIGO stage III and IV) disease, where treatment is rarely curative. Despite the addition of anti-angiogenic agents (1,2) and PARP inhibitor therapy (3), the majority of patients with advanced disease relapse within 24 months of completion of first-line chemotherapy. By contrast, the small proportion (10 - 15%) patients who present with early disease (stage I and II) have much better prognosis and are frequently cured with surgery and platinum-based chemotherapy alone (4).

HGSC is marked by near-universal *TP53* mutation (5,6) and widespread copy number change: indeed, HGSC is the archetypal C class, copy number-driven malignancy (7). The most commonly recurrently amplified genes are *MYC, MECOM, PIK3CA* and *CCNE1* (6), the latter, in particular, being associated with poor response to platinum-based chemotherapy (8). Recurrent deletions in *RB1* are seen as well as disruption to *NF1* and *PTEN* through structural variants (9). Although the genome of HGSC is highly complex, we recently described copy number signatures, recurrent patterns of genome-wide copy number change that were prognostic and were significantly associated with specific driver mutational processes (10).

Beyond *TP53*, classic driver oncogenic mutations are rare (6,9). Defective homology-mediated DNA repair mechanisms are believed to be present in approximately 50% of newly-diagnosed HGSC cases (6,11), most commonly driven by germline or somatic mutations in *BRCA1* or *BRCA2*, which are associated with improved prognosis (12) and response to PARP inhibition (13). Recent data also indicate that structural variants at the *BRCA1/2* loci are another common source of homologous repair deficiency in high grade serous ovarian carcinoma (14).

HGSC arises from the fimbriae of the distal fallopian tube, evolving from p53 signatures (cytologically normal cells with mutant *TP53*), via serous intra-epithelial carcinomas (STIC) to invasive carcinomas (15) that readily metastasise to the ovary and throughout the peritoneal cavity. However, the large studies that defined the genomic landscape of HGSC, including those from The Cancer Genome Atlas consortium (TCGA) (6), the Australian Ovarian Cancer Study (16) and the International Cancer Genome Consortium (ICGC) (9), all analysed samples from patients almost exclusively with stage III or IV disease and there is little information about the genomics of early stage HGSC. It is unclear whether early stage HGSC represents a distinct subset that fails to metastasise or whether these cases are genomically similar to late stage disease but identified essentially by chance before metastasising. To address this, we have undertaken genomic analysis, including shallow whole genome sequencing and deep sequencing of a target gene panel, of a cohort of early stage HGSC patients identified at three large UK centres, with comparison to late stage samples from the BriTROC-1 study.

## Patients and Methods

### Study conduct, survival analyses and patient samples

Details of the BriTROC-1 study have been reported previously (10,17). All other samples were identified and obtained from the pathology archives of participating hospitals by specialist gynaecological pathologists (BK, NS, JMcD) and utilised under the auspices and ethical approval of the Imperial College Healthcare Tissue Bank (HTA licence 12275, Research Ethics Committee number 17/WA/0161, Project ID R18060). All patients were identified through routine clinical practice rather than risk-reducing surgery. Overall survival was calculated from the date of diagnosis to the date of death or the last known clinical assessment. All cases underwent pathological review (CS, JMcD).

### Sequencing

Details of the sequencing of BriTROC-1 samples are given elsewhere (10). For new samples, DNA was extracted from 10 × 10 μm sections using QIAmp DNA FFPE Tissue Kit (Qiagen, UK) according to the manufacturer’s protocol. 50-200ng was sheared with a Covaris LE220 focused-ultrasonicator (Covaris, Woburn, MA) to produce 100-200bp fragments. Libraries were generated using SureSelect XT standard protocol (Agilent Technologies, Santa Clara, CA) for low-input and FFPE samples. Analysis of *PTEN, KRAS, RB1, BRCA2, RAD51B, FANCM, PALB2, RAD51D, TP53, RAD51C, BRIP1, CDK12, NF1, BRCA1, BARD1, PIK3CA* was performed using a custom Ampliseq panel on HiSeq4000 System (Illumina, Cambridge, UK), using paired-end 125 bp protocols. The mean coverage was >7000×. Shallow whole genome sequencing (sWGS) was performed on the HiSeq 4000 150 pair end (Illumina Cambridge, UK) platform, using 250-300 ng input DNA according to the manufacturer’s instructions. The minimum number of reads per sample was set at 5-10 million (mean coverage of 0.1×). Using our previous calculations (https://gmacintyre.shinyapps.io/sWGS_power/), 10 million reads with a bin size of 30kb had 80% power (alpha 0.01) to detect CN change +/− 2 at 30% purity assuming ploidy of 2.

### Somatic mutations calling from Ampliseq panel

Ampliseq FASTQ files were trimmed for adapters and aligned to reference human genome hg19 using Burrows-Wheeler Alignment (BWA-MEM) (18) and pre-processed using samtools and picard to generate sorted BAM files (19). Somatic mutations were called using mutect2 (GATK4.1.4.1) (20), Varscan2 (version 2.4.2) (21), Strelka2 (version 1.0.14) (22) and HaplotypeCaller (23) pipelines for single nucleotide variations (SNVs) and small insertions and deletions (Indels) on tumour-only BAM files, using default parameters. Mutations were annotated using Variant Effect Predictor (VEP) (version 1.5.3) (24). Somatic mutations were filtered by clinical significance with only “pathogenic” and “likely_pathogenic” mutations retained for downstream analysis. Only variants identified by at least two variant-calling algorithms were used for downstream analyses.

### Absolute copy number and copy number signature calling

sWGS reads were aligned to reference human genome hg19. Relative copy numbers were obtained for predefined 30kb bins using QDNASeqmod packages (25). We obtained absolute copy numbers using the sWGS-absoluteCN (swgs) pipeline - full details are given in Supplementary Information. Focal amplifications and deletions were defined according to the COSMIC definitions (see https://cancer.sanger.ac.uk/cosmic/help/cnv/overview): amplification was defined as total copy number ≥5 if average ploidy ≤2.7, or ≥9 if average ploidy >2.7. Loss was defined as total copy number 0 if average ploidy ≤2.7 or (ploidy minus 2.7) if average ploidy >2.7. Gain in Figure 3C was defined as total copy number >2.5 but <5.0. Copy number signatures were calculated using the R scripts as previously published (10).

### Copy number signature comparison

To model the presence or absence of signatures, a fixed effects Bernoulli model was used with an intercept and a coefficient for the change between early and late stage samples. The presence of a signature j in sample i is modelled by a Bernoulli with probability θij, where θ = x^⊤^β. x has two rows – for the intercept and the difference between the groups – and as many rows as samples. β has two rows and as many columns as the number of signatures (d = 7). The change in the differential abundance of non-zero exposures has been modelled similarly. We used a multivariate normal model based on the isometric log ratio (ILR)-transformed exposures (also called a logistic-normal model in the literature). The ILR transformation maps a d-dimensional compositional vector (the exposures) to a d−1 dimensional vector of real values. To account for absent signatures, the transformation (and subsequently the model) only used the subset of signatures that are present in each sample. Covariates are the same as those in Bernoulli model, but this time θ = x⊤β represents the ILR-transformed probabilities. Therefore, it has 6 columns instead of 7, and each row can be transformed back to a 7-dimensional vector of probabilities with the inverse ILR transformation. β will continue to have two rows, but only 6 columns, which indicate changes in log-ratios of signature exposures. The R package TMB (26) was used for inference. The model has been written in C++. The code is publicly available at https://github.com/lm687/Cheng_exposure_analysis and a full description of the analysis is given in Supplementary Methods.

### Unsupervised clustering of patients using signature exposures

Hierarchical clustering of the exposure vectors of samples (early stage and late stage) used in the survival analysis was performed using the NbClust (27) package in R. A Cox proportional hazards model was fitted using the cluster labels as covariates, stratified by stage (early vs late), using the R packages survival (28) and survminer (https://rpkgs.datanovia.com/survminer/index.html).

### Statistical analyses

Unless otherwise stated above, statistical analyses were performed using Prism (v9.0.3, GraphPad, CA).

## Results

### Patients and samples

We initially identified 54 patients with early stage (defined as stage IA, IB, IC and IIA using the FIGO classification at the time of diagnosis) ovarian high grade serous carcinoma from the pathology archives of three large UK gynaecological cancer centres (Imperial College Healthcare, University College London and Barts Health NHS Trusts). A summary of the workflow is shown in **Fig. 1** and clinical details are given in **Table S1**. Following pathology review, 21 samples from 13 patients were excluded, whilst two samples from one patient failed DNA extraction. Additionally, we identified a further cohort of five early stage patients recruited into the BriTROC-1 study (17), giving a total early stage population of 45. The comparison late stage cohort consisted of 51 randomly selected patients with stage IIIC/IV disease recruited into the BriTROC-1 study. The median age at diagnosis for early and late stage cohorts did not differ significantly (early 61.3 years, range 40-84; late 62.3 years, range 34-76) but overall survival was, as expected, significantly longer in the early stage cohort than for the late stage (Hazard Ratio 0.11, 95%CI 0.06-0.22) (**Fig. S1**).

**Figure 1.**
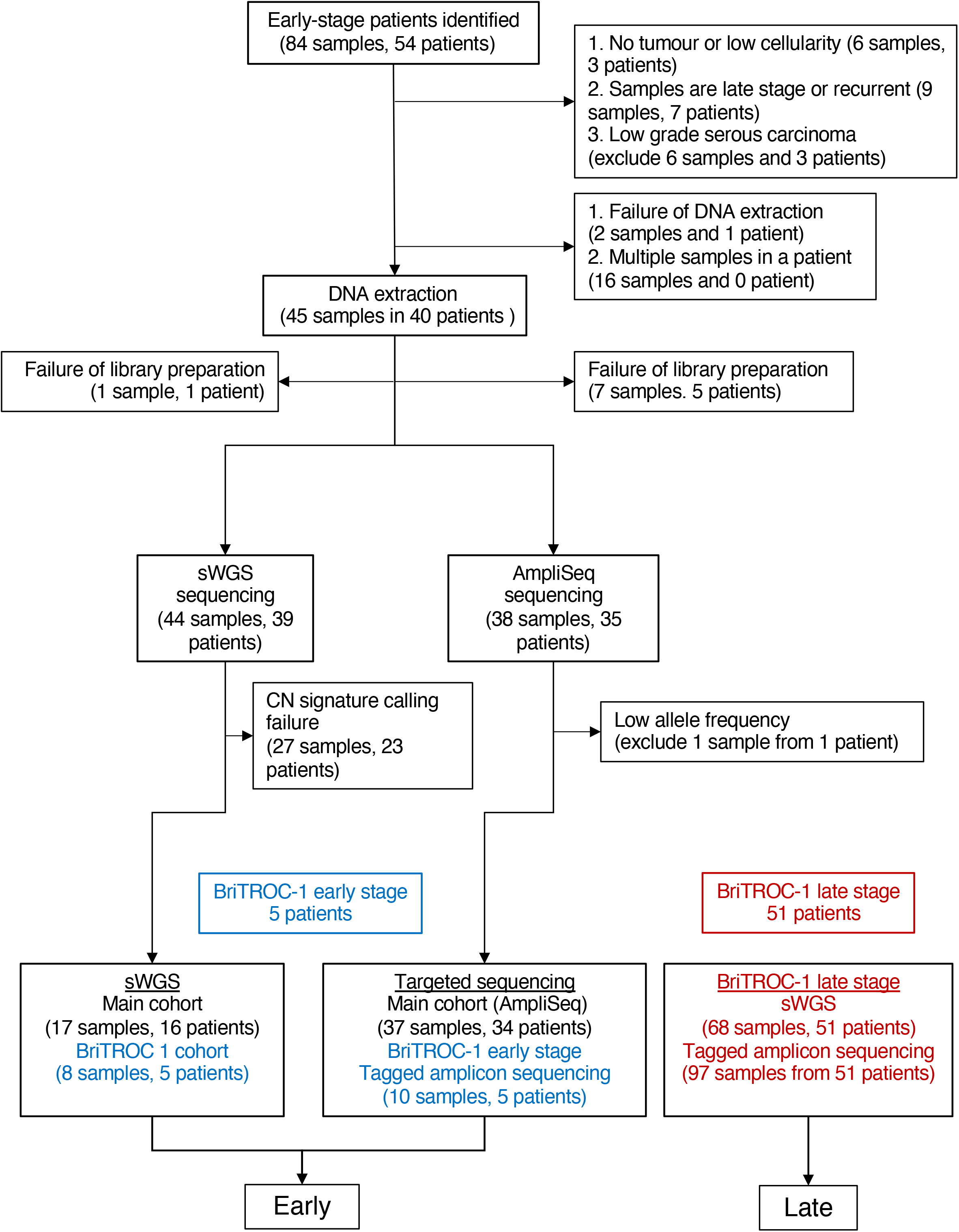
REMARK diagram for early stage and late stage cohorts.

### Mutational landscape of early stage and late stage cohorts

Using targeted next generation sequencing, we analysed short variants (SNV, indels) in both cohorts (**Table S2**). Mutations in *TP53* were near-universal (100% late stage: 97/97 samples, 51/51 patients. 92% early stage: 45/49 samples, 35/39 patients. **Fig. 2A**). The four early stage samples in which *TP53* mutations were not identified underwent pathology re-review; all were confirmed to be HGSC (**Fig. S2**). The frequency of four key *TP53* hotspot mutations (R175, R273, R248, Y220) was significantly greater in the early stage cohort compared to late (**Fig 2B, C**. Fisher’s exact test; p=0.0075). There was no difference in the rates of mutations in the other analysed genes (**Fig. 2A**). Specifically, the rates of pathological mutations in *BRCA1* and *BRCA2* were 10% and 2% respectively in the early stage and 14% and 2% in the late stage.

**Figure 2.**
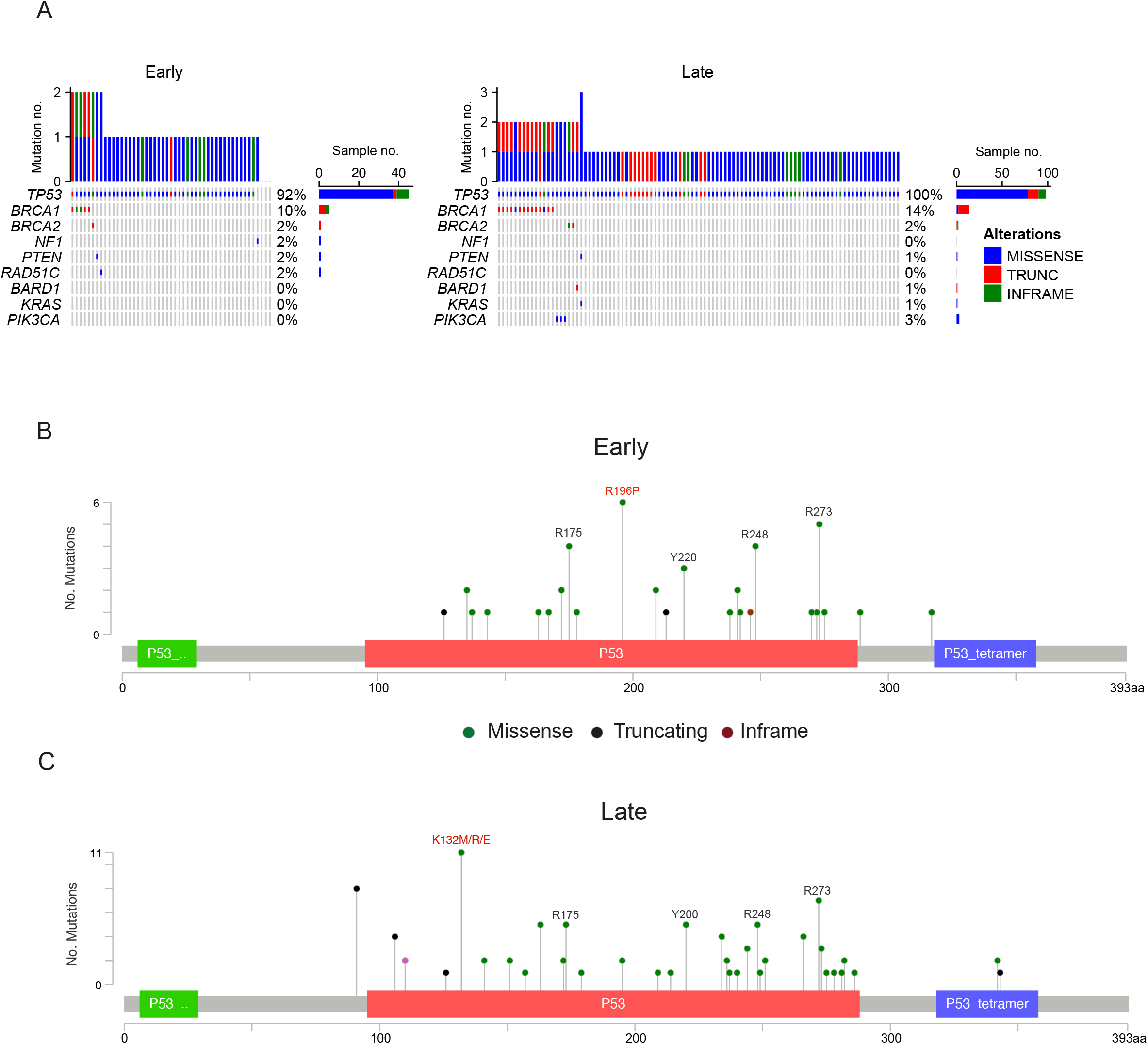
Mutational landscape of early stage and late stage cohorts. A. Somatic mutation rates in early stage and late stage cohorts. The upper plot shows the frequency of mutation for each tumor sample. B. Gene mutation mapper plot of *TP53* in early stage cohort and (C) late stage cohort. Key hotspot residues are marked. The commonest residue mutations in each cohort are marked in red

### Focal amplifications and deletions in early stage and late stage cohorts

We used shallow whole genome sequencing to analyse genome-wide absolute copy number. There was no statistically significant difference in purity between the cohorts (**Fig. 3A**), but median ploidy was significantly greater in late stage samples compared to early (**Fig. 3B** Median early 2.0; median late 3.1; p<0.0001). Global copy number gains/losses are shown in **Fig. 3C**-there were generally more gains and amplifications in late stage samples, and regions of CN loss in the early stage samples, in keeping with the differential ploidy between the cohorts. Although there were several regions of differential gain in the late stage cohort (e.g. chromosome 4, 6, 9, 11,12) and losses in the early stage cohort (e.g. chromosome 4, 9, 12, 17), we found no significant differences in rates of focal amplification and deletion of 16 of 17 genes that are frequently altered in HGSC (6,9) (**Fig 3D, Fig. S4**). The commonest amplifications were in *MYC* (24% in the early stage and 21% in the late stage) and *MECOM* (20% in the early stage and 15% in the late stage). The only gene with differential CN change was *PTEN*, which was amplified in 4/25 early samples compared to only 1/58 late stage (p<0.05) - the rates of *PTEN* deletion, however, did not differ between the cohorts.

**Figure 3.**
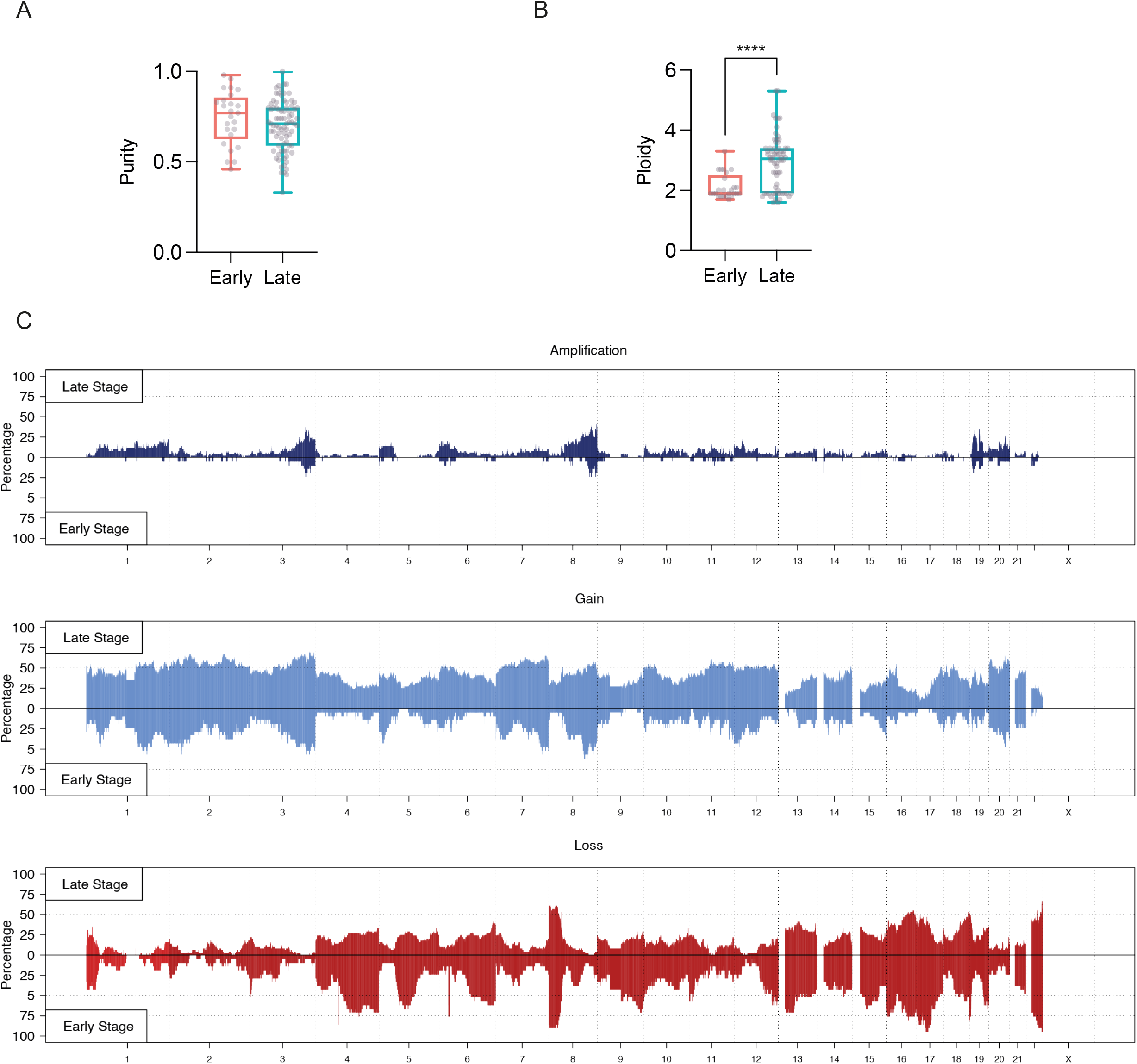
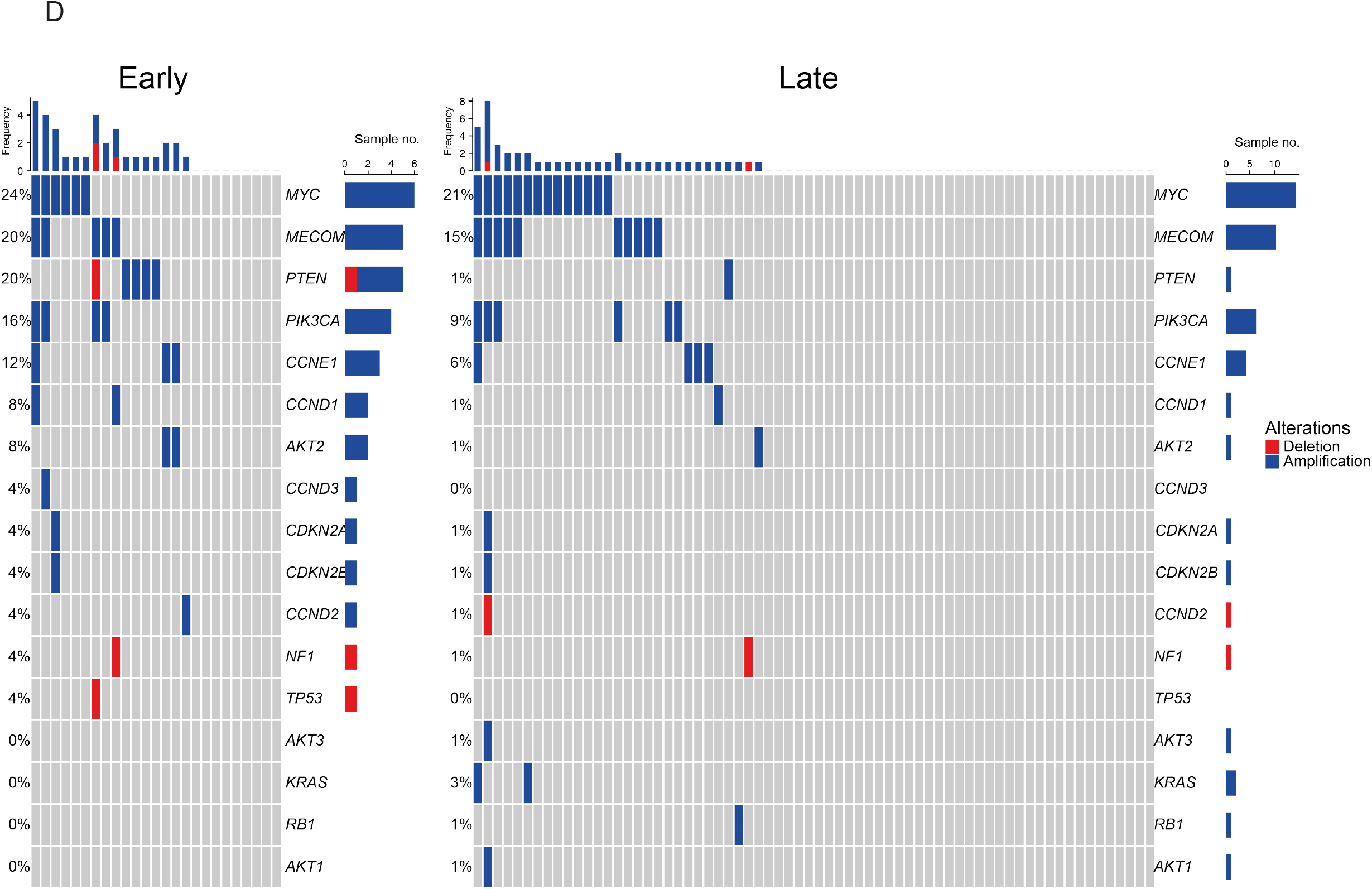
Focal gene amplifications and deletions in early stage and late stage cohorts. (A) Purity comparison of early stage and late stage cohorts. (B) Ploidy comparison of early stage and late stage cohorts; Mann Whitney test. ****; p<0.0001 (C) Global copy number amplification, gains and losses in early stage and late stage cohorts. (D) Absolute copy number estimation in 17 genes of interest, determined by sWGS.

### Copy number signatures in early stage and late stage cohorts

Next, we assessed the distribution of the six specific CN features - segment length, segment copy number, number of breakpoints per chromosome arm, number of breakpoints per 10Mb, copy number change point and length of chains of oscillating copy number - that we previously used to derive copy number signatures in HGSC genomes (**Fig. 4A**). Distributions normalised per sample (**Fig. 4B, Fig. S5**) showed that late stage HGSC genomes were significantly more likely to have 2 or more breakpoints per 10Mb and more than 4 breakpoints per chromosome arm than early stage. In addition, late stage genomes had higher copy number change point, segment copy number and smaller segment lengths than the early stage genomes (**Fig. 4C, Fig. S5**). We then generated CN signature exposures (**Fig. 5A, B**) for both cohorts and used a fixed-effects (Bernoulli) analysis to model the presence or absence of signatures and a fixed-effects multivariate normal distribution model, based on isometric log ratio (ILR)-transformation, to compare the two cohorts (see Supplementary Methods). Overall, in the ILR analysis, there was a significant difference between cohorts (generalised Wald test; p=0.0015), with greater signature 3 exposure in the early stage cohort and more signature 4 in the late stage cohort (**Fig. 5C**).

**Figure 4.**
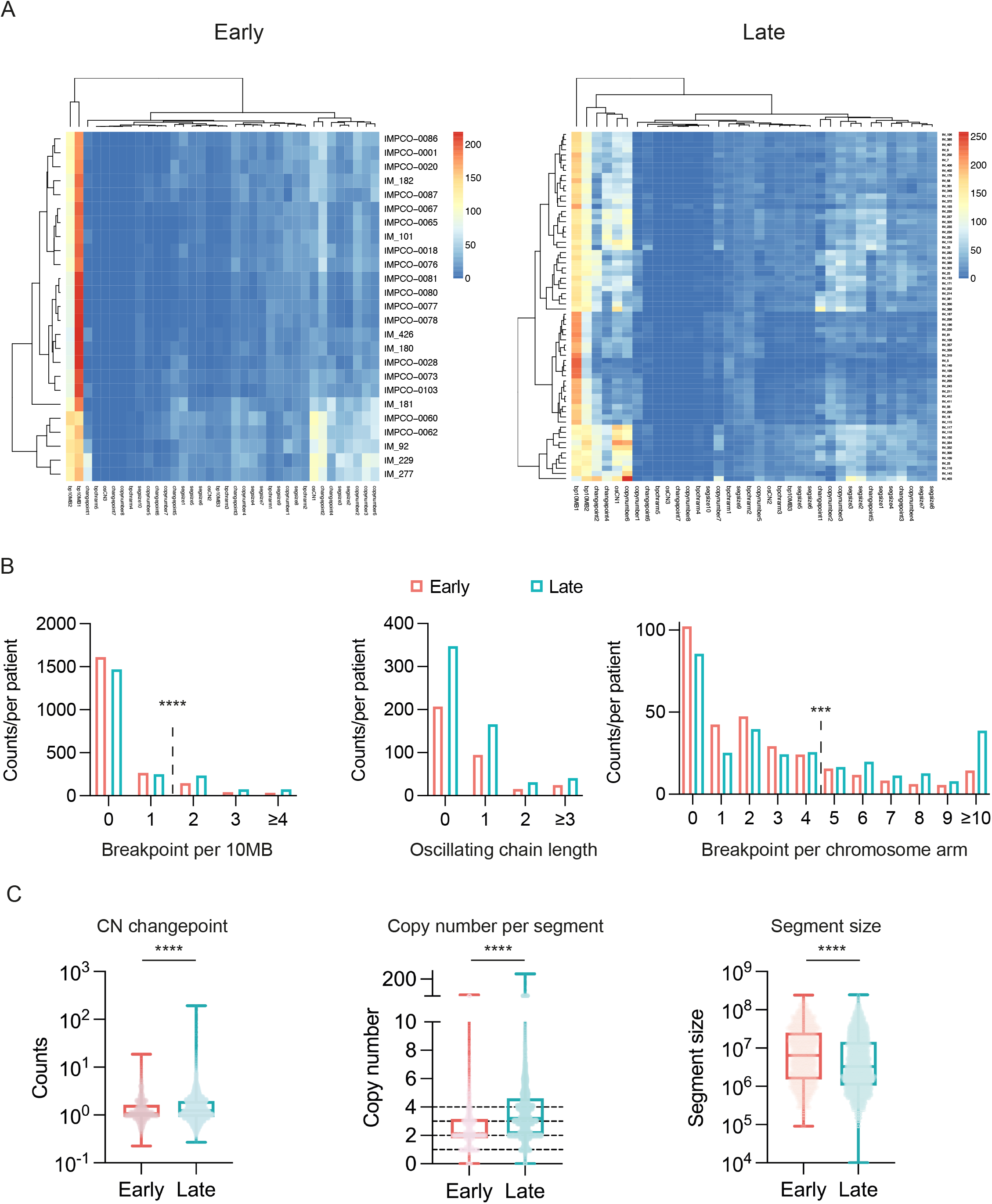
Components and features distribution in early stage and late stage cohorts. (A) Tumour-by-component matrix distribution of 36 copy number components in early stage (left) and late (right) stage cohorts. (B) Counts per patient in copy number features - Breakpoint per 10MB, Oscillating chain length and Breakpoint per chromosome arm; Fisher’s exact test (C) Counts in copy number features - copy number changepoint per segment, absolute copy number per segment, segment size); Kolmogorov-Smirnov test ***; p<0.001 ****; p<0.0001

**Figure 5.**
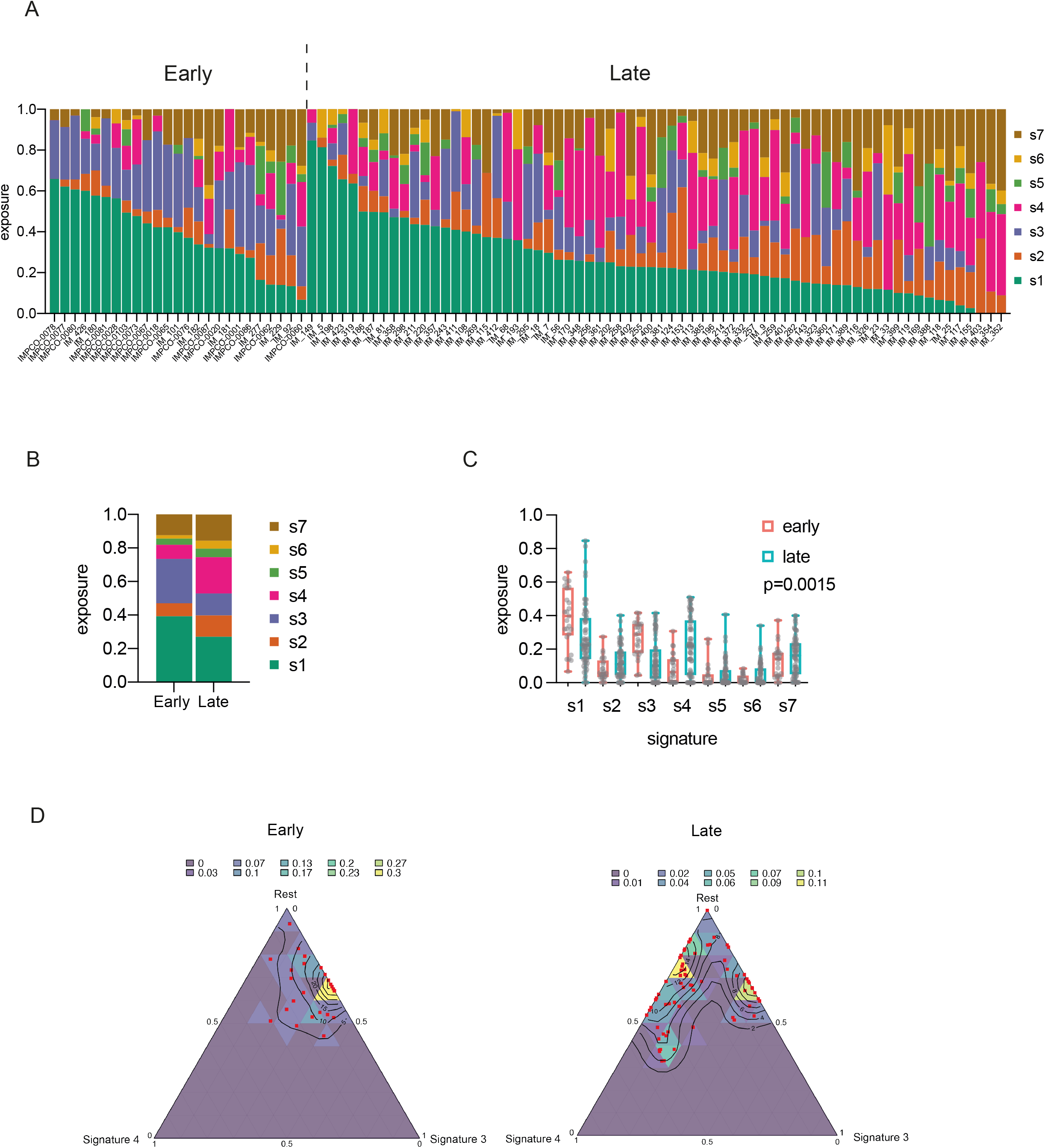
Copy number signatures in early stage and late stage cohorts. (A) Copy number signature exposures in early stage and late stage cohorts. (B) Mean signature exposure proportions across the early stage and late stage cohorts. Note that signature exposures sum to 1 in each patient. (C) Comparison of signature exposures across early stage and late stage cohorts; Wald test (D) Simplex plots representing exposures for CN signature 3 (right axis), signature 4 (bottom axis) and the rest of the signatures (1 - S3 - S4) combined (left axis) in early (left) and late (right) stage cohorts. Each red dot represents a single sample and the contours represent the density of observed samples.

We then visualised CN signature exposures using simplex plots (**Fig. 5D**) comparing signature 3 (S3), signature 4 (S4) and all other signatures (1-S3-S4). In the early cohort, the sample observations (red dots) cluster towards the top of the right side of the simplex, in keeping with low or zero S4 exposure. For the late group, although some of the observations remain in the same place, many are located towards the left of the simplex, indicating that they have non-zero exposure to S4, with a relative decrease in the amount of S3. The relative contribution of the other signatures does not change between early and late cohorts - the distance from the observations to the top apex of the plot remains similar. Together, this suggests that overall S3 decreases in intensity and S4 increases in intensity in the late stage samples, whilst the rest of the signatures remain approximately constant. However, the observations that remain in the same place in the simplex suggest that a subset of late stage samples have genomic features more reminiscent of early stage.

Finally, we performed unsupervised hierarchical clustering of the copy number signature exposures across both cohorts, and identified three clusters (**Fig. 6A**). Cluster 1 had the highest exposure to CN signature 3, cluster 2 was dominated by genomes with high signature 1 exposure, whilst cluster 3 showed high signature 4 exposure. There was a significant difference in sample distribution between clusters (p<0.0001, Chi-squared) with nearly all early stage samples in clusters 1 and 2, whilst late stage samples were spread across all three clusters, with the majority in cluster 3 (**Fig. 6B**). The clusters were prognostic, with a significant trend for reduced survival across the clusters (**Fig. 6C**), which remained significant in multivariate analysis stratified by stage (early vs late; **Fig. 6D**). We also quantified ploidy in the late stage samples by cluster and found highly significant differences (**Fig. 6E**), with median ploidy of 1.9, 2.2 and 3.3 in clusters 1, 2 and 3 respectively, suggesting that a subset of late stage samples (those in cluster 1) retain genomic features of early stage disease, whilst the majority (clusters 2 and 3) evolve, with emergence of increased ploidy.

**Figure 6.**
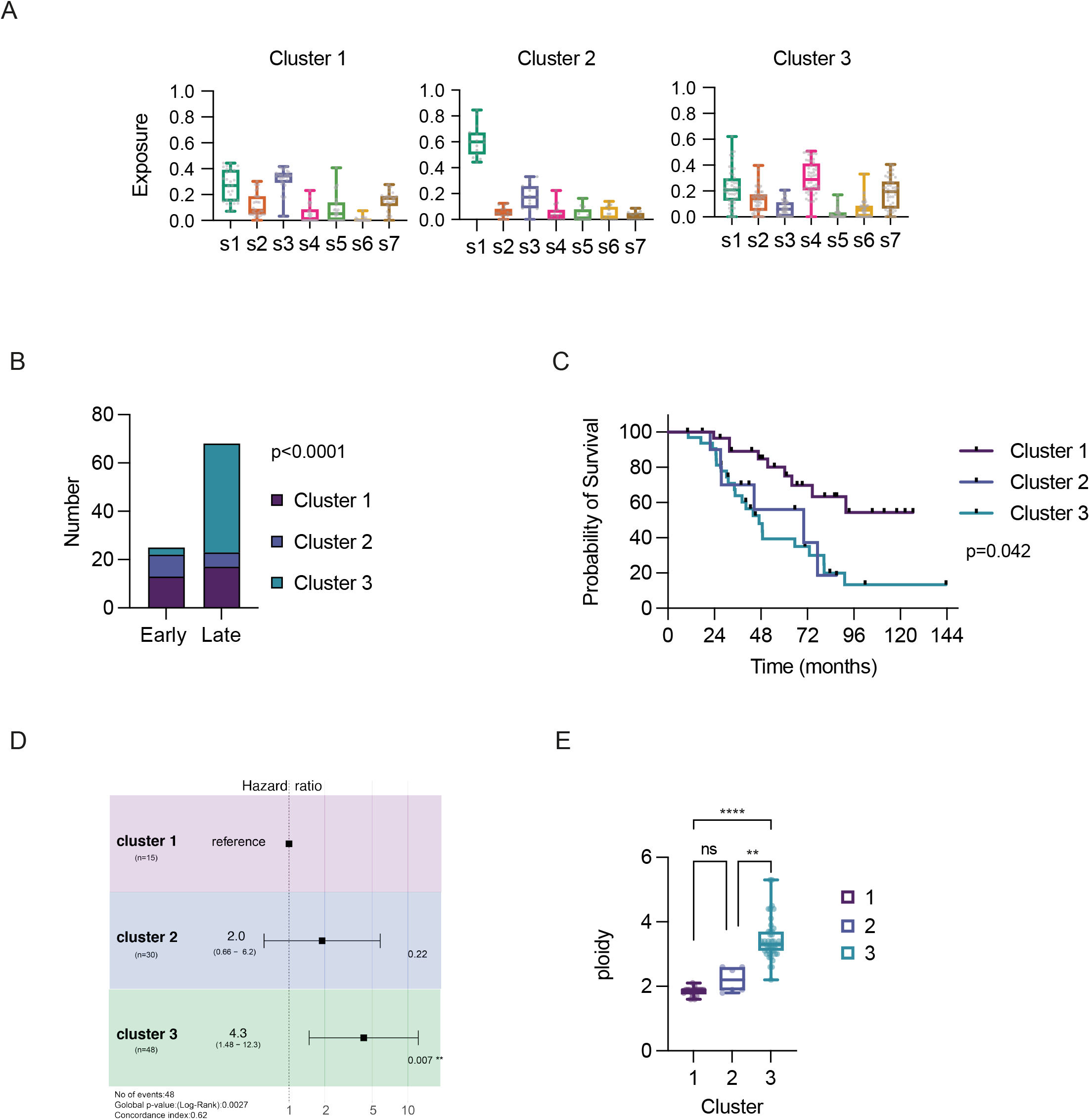
Relationship between signature exposures and clinical factors. (A) Unsupervised hierarchical clustering in combined early stage and late stage cohorts. Distributions of copy number signature exposures in three clusters (B) Early and late stage samples by cluster; Chi-squared (C) Overall survival by cluster; Log-rank (Mantel-Cox) analysis (D) Ploidy distribution of late stage samples in three clusters. One-way ANOVA with multiple comparisons. **;p<0.01, ****p<0.0001

## Discussion

The large majority of patients with HGSC have advanced disease at the time of diagnosis, reflecting the ease with which HGSC disseminates throughout the peritoneal cavity. An important question for early stage HGSC is whether these tumours have discrete characteristics that reduce the likelihood of metastases or whether these tumours were identified fortuitously through earlier development of symptoms.

We used targeted next generation sequencing and shallow whole genome sequencing in a cohort of stage I - IIA samples and compared to stage IIIC/IV samples from the BriTROC-1 study. All patients in our cohort were identified through routine clinical practice rather than being diagnosed following risk-reducing surgery, and it was noteworthy that there was no significant difference in median age at diagnosis between our two cohorts. *TP53* mutations were near-universal in both early- and late-stage cases as expected (5). All samples underwent pathology review as differentiating high grade from low grade serous carcinoma can be challenging, and six samples were re-classified as low grade following review. Four other samples that were classified as HGSC by pathologists were *TP53* wild type, possibly due to a combination of low tumour cellularity and poor DNA quality from FFPE preservation - certainly two of these samples had copy number profiles strongly suggestive of HGSC (**Fig. S2**). Interestingly, we found that the rate missense *TP53* mutations was higher (75%) than in previous HGSC cohorts (6) and that *TP53* hotspot mutations (defined here as mutation at the four most commonly mutated codons R248, R273, R175 and Y220) were significantly more prevalent in our early stage cohort than late stage (**Fig. 2B,C**). Previous analyses of the prognostic effect of *TP53* mutation type and location in HGSC have provided contradictory results (5,29). Recently, a detailed analysis of nearly 800 HGSC cases (78 stage I/II, 709 stage III/IV) suggested no overall difference in mutation type between early and late samples, but three specific hotspot mutations G266, Y163C and R282, were associated with poorer overall survival (30). However, only one early stage patient, ES_007, had one of these mutations and was alive 109 months following diagnosis. Beyond *TP53*, we found no differences in rates of *BRCA1/2* mutations between early and late cohorts. One limitation of our study was the absence of germline DNA from the majority of our early stage cohorts - these samples were identified through pathology archives - and so we were unable to verify the germline status of the samples with pathogenic mutations. However, our *BRCA1/2* mutation results are broadly in line with previous cohorts (31) and reflect the fact that these samples were obtained from routine practice rather than risk reducing surgery.

When we compared focal copy number alterations in 17 key genes, again we found no significant difference between the two cohorts in 16 of the 17 (**Fig 3D**), although, curiously, we did find more *PTEN* amplifications in the early stage cohort: we believe that these are unlikely to be of clinical significance and are likely to reflect the overall genomic instability of HGSC. When examining global copy number change, we observed some significant changes in the proportion of cases with gain/loss at certain loci (**Fig 3C**) but there were no regions uniquely lost or gained in either cohort, suggesting that the process of dissemination is not driven by specific amplifications or deletions. These SNV and focal CNA data corroborate findings by Köbel et al that early and late stage HGSC appear identical by immunohistochemistry (32).

In addition to the absence of matched germline DNA, this project has other potential shortcomings. Our cohort is relatively small, reflecting the rarity of this patient population, which limits our statistical power. However, this still represents one of the largest collections of early stage HGSC samples that have been characterised at a genomic level. The samples were all formalin-fixed, paraffin-embedded (FFPE) and analysed up to 10 years following diagnosis. Consequently, several samples failed sequencing and CN signature calling. In addition, the BriTROC-1 patients were all recruited at relapse (17), necessitating analysis of FFPE material from the time of diagnosis for the comparator late stage cohort. We also did not perform whole exome sequencing or deep WGS, so are unable to comment upon small variants (SNV, indel) beyond our targeted panel nor on larger scale rearrangements that are prevalent in HGSC (9,14). Thus, we cannot confirm all the SV and LOH findings from a previous study that performed deep WGS on 16 early stage HGSC samples (33). However, like us, Chien et al identified near-universal *TP53* mutations and high levels of genomic instability in their cohort with few, if any, recurrent focal differences between early and late stage HGSC, and certainly no unique mutation or focal CN alterations in the early stage patients. They identified a significantly higher rate of *PRIM2* loss in early stage HGSC, which we did not observe here.

Our most striking observation was the difference in overall ploidy between early and late cases; early stage tumours were largely diploid whilst the median ploidy in late stage samples was >3. This was further reflected in copy number (CN) signature exposures, patterns of copy number change derived from analysis of six copy number features that may reflect underlying driver mutational processes (10). Comparison of CN signatures between samples and cohorts is complex because signature exposures are compositional (i.e. they sum to 1 in each sample) and are thus not independent variables: any decrease in one signature will, by definition, be mirrored by an increase in at least 1 other signature. In addition, classical methods for analysing compositional data are poor at dealing with zero proportions and many samples have zero exposure to at least one signature (**Fig. 5A**). However, using isometric-log ratio analysis of non-zero signature exposures, we found a significant difference between the cohorts overall, driven by signatures 3 (higher in early stage) and 4 (higher in late stage). The features that define signature 4 are high segment copy numbers and high copy number changes, both of which are significantly greater in the late cohort (**Fig. 4C**). In addition, the late stage genomes were more likely to have a higher number of breakpoints per 10MB and per chromosome arm and smaller segment size (**Fig. 4B, C**), suggesting evolving genomic instability. The higher copy number, greater overall ploidy and greater signature 4 exposure all suggest that a proportion of the late stage tumours have undergone whole genome duplication; certainly, CN signature 4 was significantly associated with whole genome duplication in our original analysis (10). The potential evolution of genomes was corroborated by both the simplex analysis (**Fig. 5D**) and the unsupervised clustering of CN signatures (**Fig. 6**). The simplex plots indicate that, whilst the late stage genomes overall have increased signature 4, a proportion remains ‘early-like’ with prominent CN signature 3. The clustering identified three patterns in the signatures. Clusters 1 and 2 contained most of the early stage samples, whilst the late stage samples were divided between the clusters. However, cluster 3 contained almost exclusively late stage samples and was associated both with higher ploidy and worse survival.

Whole genome duplication (WGD), duplication of a whole set of chromosomes, has been described in many solid malignancies (34) and is generally associated with poor prognosis (35). It is thought to arise from aberrant cell division (36) and potentially may mitigate the effects of mutations that would otherwise be deleterious (37). In their analysis of over 9000 cancers of multiple types, Bielski et al identified that the median ploidy of tumours of all types that had undergone WGD was 3.3, compared to 2.1 in those lacking WGD, strikingly similar to our late and early cohorts respectively. Indeed, ovarian high grade serous carcinoma had one of the highest rates of WGD at approximately 40% (34). The commonest genomic correlate of WGD was mutation in *TP53*, which usually precedes the duplication and is near-universal in HGSC, as well as *CCNE1* amplification and loss of *RB1*. Interestingly, in prostate, pancreatic and non-small cell lung cancers, but not endometrial or breast cancers, the rates of WGD were significantly greater in metastatic deposits than in primary tumours. Although the rates of *CCNE1* amplification did not differ significantly between our early and late stage cohorts, our data support the idea that WGD is associated with advanced HGSC and poorer prognosis. HGSC generally has profound levels of segregation error during cell division, an essential precursor of aneuploidy and WGD (38), and recent data suggest that WGD can emerge in *hTERT*-immortalised human fallopian tube epithelial cells with loss of wild-type p53 function in the presence of *BRCA1* mutation and *MYC* over-expression, but not with *TP53* mutation alone (39). However, one outstanding question is whether WGD *per se* promotes rapid dissemination in HGSC, or whether it is a time-related consequence and is thus more likely to be observed in late stage than early stage disease: detailed multi-site analysis of disseminated HGSC and *in vitro* models will be required as well as further analyses of early stage samples to address this question.

In summary, our results indicate that early and late stage HGSC are remarkably similar but also that there is likely to be a degree of genomic evolution from early to late stage, potentially resulting from the appearance of whole genome duplication, which is associated with poor outcome. However, our data, reinforced by the striking difference in overall survival in our cohorts, highlight once again the importance of strategies that will allow early detection of HGSC.

## Supporting information

Supplementary Tabl 2

## Acknowledgements

This work was funded by an Imperial/China Scholarship Council scholarship to ZC, the NIHR Imperial Biomedical Research Centre (grant number P77646), Ovarian Cancer Action, the Wellcome Trust (grant number RG92770) and Cancer Research UK (grant numbers A15973, A15601, A18072, A17197, A19274 and A19694). Infrastructure support was provided by Experimental Cancer Medicine Centres at participating sites, the Cancer Research UK Imperial Centre and the Imperial College Healthcare Tissue Bank.

## Author contributions

The authors have declared no conflicts of interest.

Study design: IAMcN, FM, JDB

Sample acquisition: ML, LT, JK, DE, BK, NS, JMcD

Pathological assessment: BK, NS, CS, JMcD, DE

Data acquisition: ZC, HM, PS, GG, DE

Data analysis: ZC, HM, PS, LM, TG, TB, AP, GM

Manuscript preparation: ZC, IAMcN

All authors reviewed the manuscript before submission

