## Supplementary methods and figures for "The genomic landscape of early stage ovarian high grade serous carcinoma"

### Supplementary information

#### BriTROC authors:

R.M. Glasspool, Beatson West of Scotland Cancer Centre
C. Gourley, University of Edinburgh
R. Kennedy, Queen's University Belfast
R. Naik, Gateshead Health NHS Foundation Trust
G. Hall, University of Leeds
R. Edmondson, University of Manchester
A. Clamp, Christie Hospital NHS Trust
S. Sundar, University of Birmingham
A. Walter, University Hospitals Bristol NHS Trust
M. Hall, East and North Hertfordshire NHS Trust
H. Gabra, Imperial College London
C. Fotopoulou, Imperial College Healthcare NHS Trust
E. Brockbank, Barts Health NHS Trust
A. Montes, Guys and St Thomas' Hospital NHS Trust
All in the UK

### Supplementary Methods

#### Absolute copy number

The sWGS-absoluteCN (swgs) pipeline implements a method for generating absolute copy number profiles from shallow whole genome sequencing data utilising a multi-stage fitting process to generate a set of read depth-normalised absolute copy number profiles. Copy number fits are generated in read count space using a modified implementation of QDNAseq (25) on full read depth shallow whole genome BAM files. After which, a grid search of ploidy and purity values is performed to generate a matrix of potential absolute copy number profiles. The fitting of absolute copy number profiles includes a fit error function and a variant allele fraction anchoring process which assist in the selection of an optimal copy number fit. Minimisation of the clonality error function, expected-to-experimental TP53 distance, and manual review of fits allows for the selection of an optimal absolute copy number profile for a given sample. Selected fits are subject to a calculation of power to detect copy number alterations in which the read depth of a given sample is assessed against its selected ploidy-purity combination. This process both determines if 1) a sample has a sufficient number of reads to support the selected ploidy-purity combination and 2) an optimal target number of reads to perform sample-specific read down sampling. This process normalises the read depth between samples in a ploidy and purity dependent manner so that read variance across segments is consistent, while excluding samples which are not supported by a sufficient number of reads. Samples passing all filtering criteria then undergo read down sampling to the specified target number of reads determined by the previous steps and absolute copy number profiles fitted at the ploidy-purity combination selected prior. We also used ACE (40) to calculate the ploidy and purity of each sample as a validation of swgs pipeline with the settings “penalty=0.5” and “penploidy=0.5” and bin size=30kb.

#### Comparison of copy number signature exposures in early and late stage patients

##### 1. Analysis of zero exposures

Signature 1 and signature 3 are always present in early samples, but sometimes absent in late samples. The change is particularly noteworthy for s3, which goes from being present in all samples of the early group to being absent in 24% of samples of the late group.

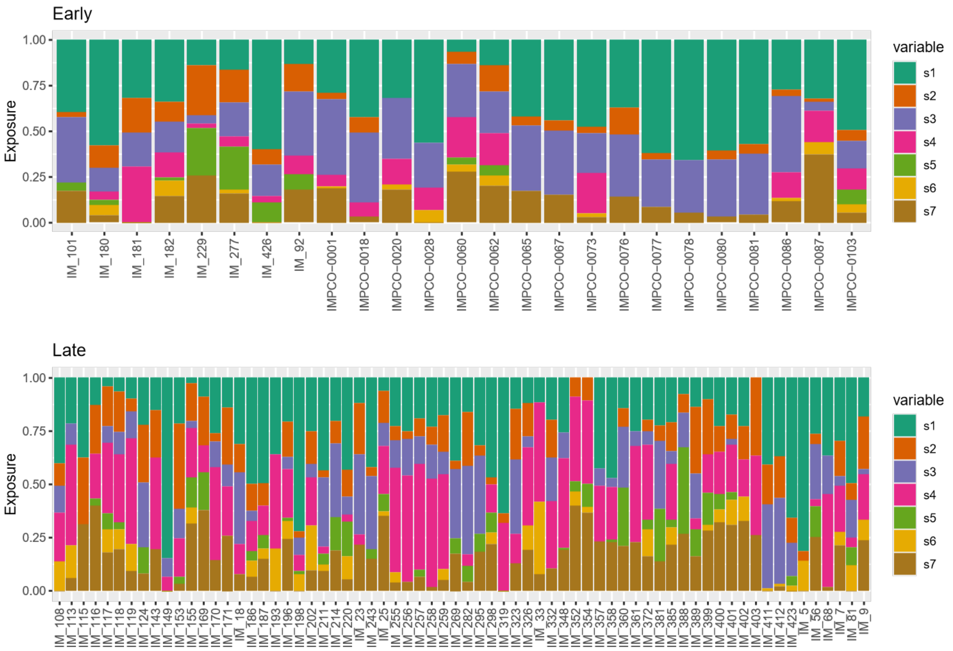


Figure 1: Exposures in each group

| | early | late |
|---|---|---|
| s1 | 0.00 | 0.04 |
| s2 | 0.12 | 0.13 |
| s3 | 0.00 | 0.24 |
| s4 | 0.32 | 0.19 |
| s5 | 0.60 | 0.46 |
| s6 | 0.52 | 0.49 |
| s7 | 0.12 | 0.13 |

Table 1: Fraction of zero exposures for each group/signature combination

s1 and s3 are removed from the following analysis of zero exposures. The model treats independently each signature, so excluding them is not problematic.

The only potential result that we are not reporting in this analysis is s3. Whereas in s1, only three samples have a zero exposure in the late group, there are 16 samples with inactive s3. As the model is not suitable because the two groups show a perfect separation, we need another way of assessing whether s3 is less present in the *late disease* group. Using a generalised Wald test for the global presence and absence of signatures, the results are not statistically significant. However, from the estimates of the slopes we can see that we have good indication that there are fewer zero s4 exposures in the late cohort.

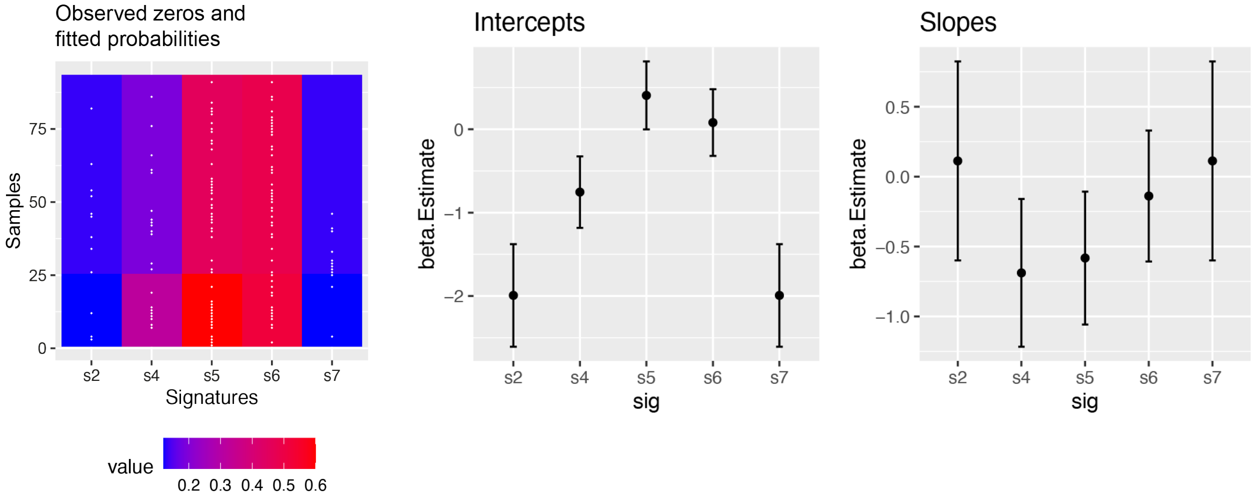


Figure 2: Results for the Bernoulli model of present/absent signatures. (Left) Observed zeros (white dots) and fitted probabilities of a zero exposure (colour-coded). (Right) estimates and standard errors for the intercepts, and estimates and standard errors for the slope (i.e. change between the two groups).

#### 2. Partial ILR analysis

Here, we analyse non-zero exposures.

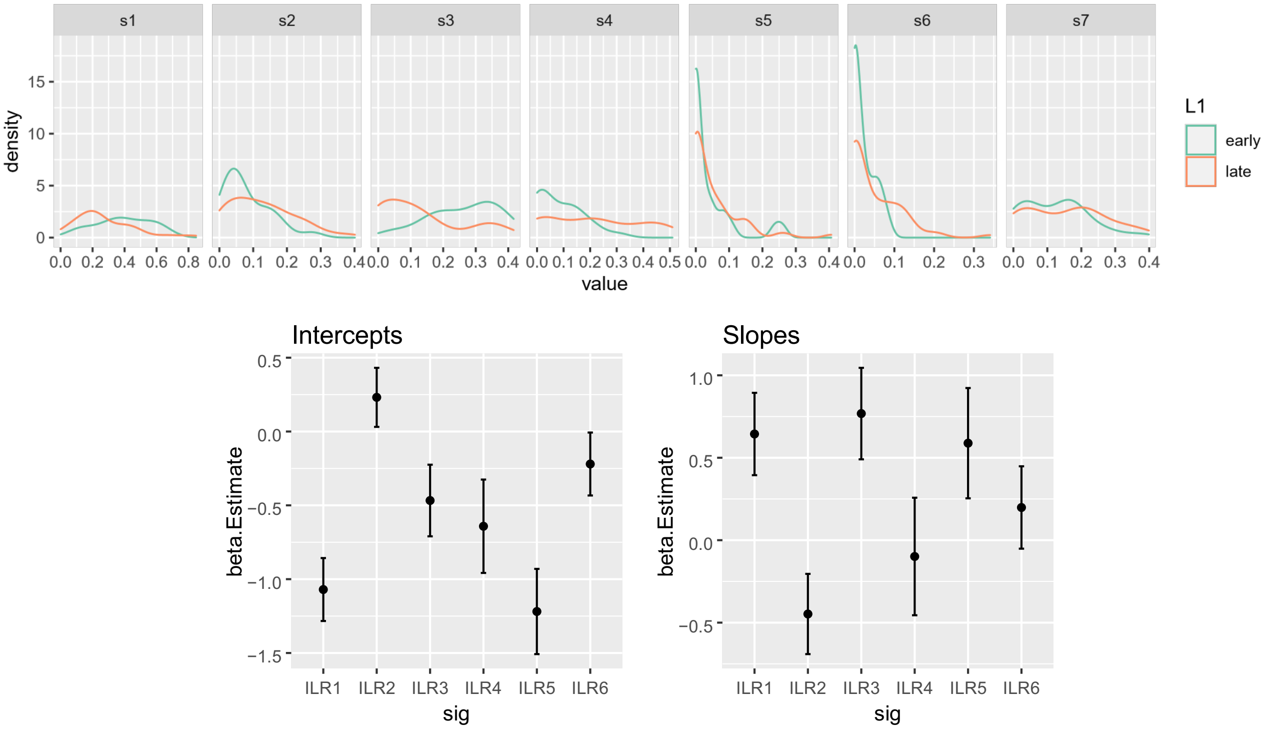


Figure 3: Partial ILR analysis for the analysis of nonzero exposures. (a) Density plots for each signature, in each group. (b) Estimates and standard errors for the intercepts. (c) estimates and standard errors for the slope (i.e. the change between the two groups).

The density plots and the slope coefficients can be summarised as follows:

- There is a relative increase of s2 in the late sample cohort (see ILR1) with respect to s1.
- There is a relative decrease of s3 in the late sample cohort (see ILR2)
- There is a relative increase of s4 in the late sample cohort (see ILR3)

A generalised Wald test indicates that the difference between the two groups is statistically significant ($p$ = 0.0015).

#### 3. Summary

In summary, we have used a fixed-effects Bernoulli to model the presence/absence of signatures, and a fixed-effects multivariate normal distribution on the transformed data to model subsets of nonzero exposures. The transformation used is a variant of the ILR transformation which is used in Compositional Data Analysis.

The findings indicate that s3 has higher exposures in the early group, both analysing zero and nonzero exposures. S4 behaves in the opposite direction.

#### Supplementary Figures and Legends

**Supplementary Table 1.** Clinical information on early and late stage cohorts

**Supplementary Table 2.** Results of AmpliSeq and Tagged-amplicon sequencing from both cohorts.

**Supplementary Fig 1**. Clinical information on early stage and late stage cohorts.
(A) Diagnosis age. Median 61.3 year (early stage), 62.3 years (late stage). p=NS
(B) Overall survival. Median 60.3 months for late stage, and not reached for early stage. Log-rank Hazard Ratio 0.11 (95%CI 0.06-0.2), $p<0.0001$ (Log-rank Mantel Cox).

**Supplementary Figure 2.** Relative copy number profiles of early stage four samples without *TP53* mutations.

**Supplementary Fig 3**. Ploidy and purity comparison between swgs and ACE pipelines. Spearman correlation test, $p<0.05$.

**Supplementary Fig 4.** Absolute copy number (y-axis) of 17 individual genes in early and late stage samples. *; $p<0.05$, **;$p<0.01$, ***;$p<0.001$, ****;$p<0.0001$

**Supplementary Fig 5.** Raw data on copy number feature distributions in early and late stage HGSC

Figure S1

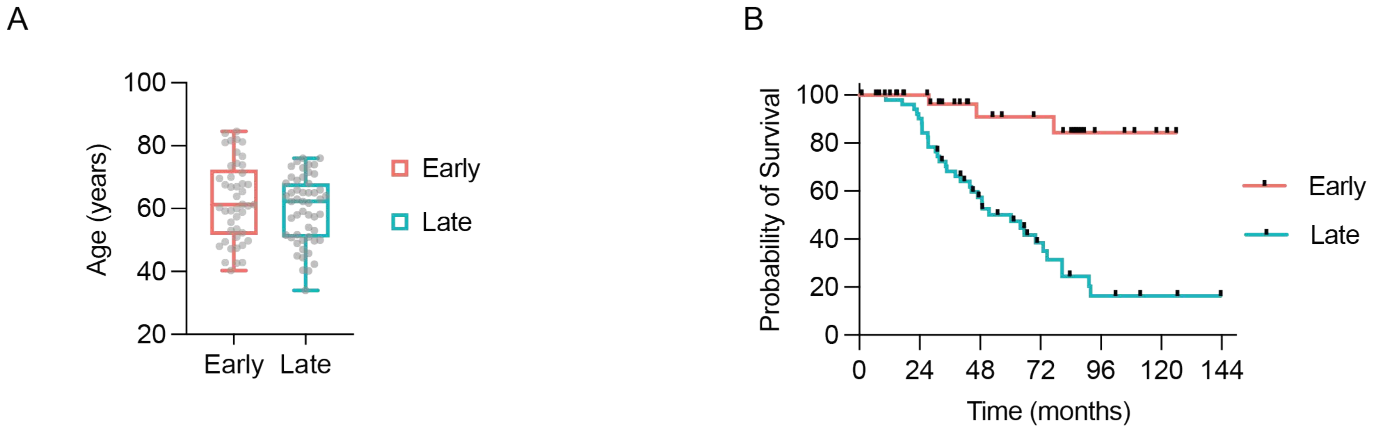


Figure S2

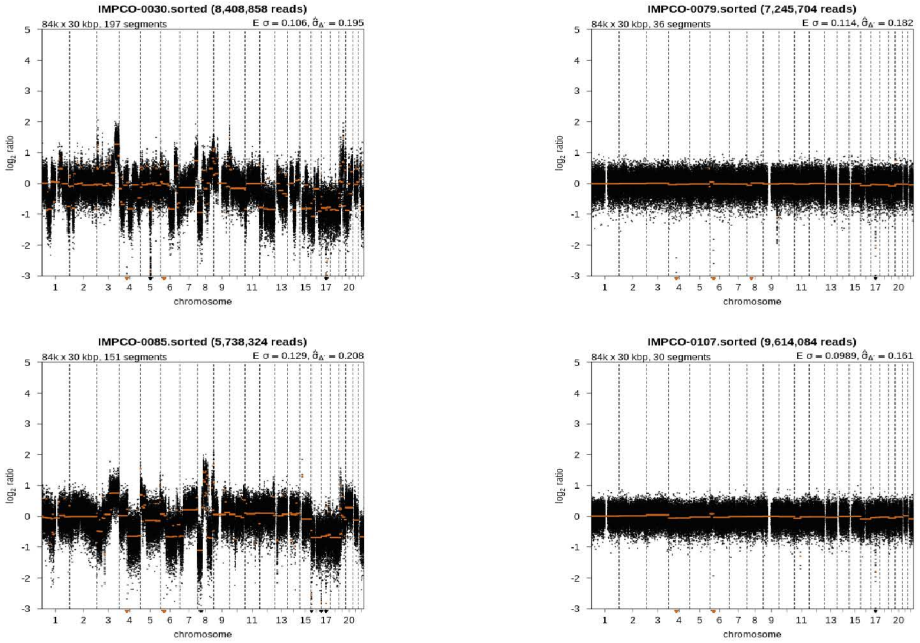


Figure S3

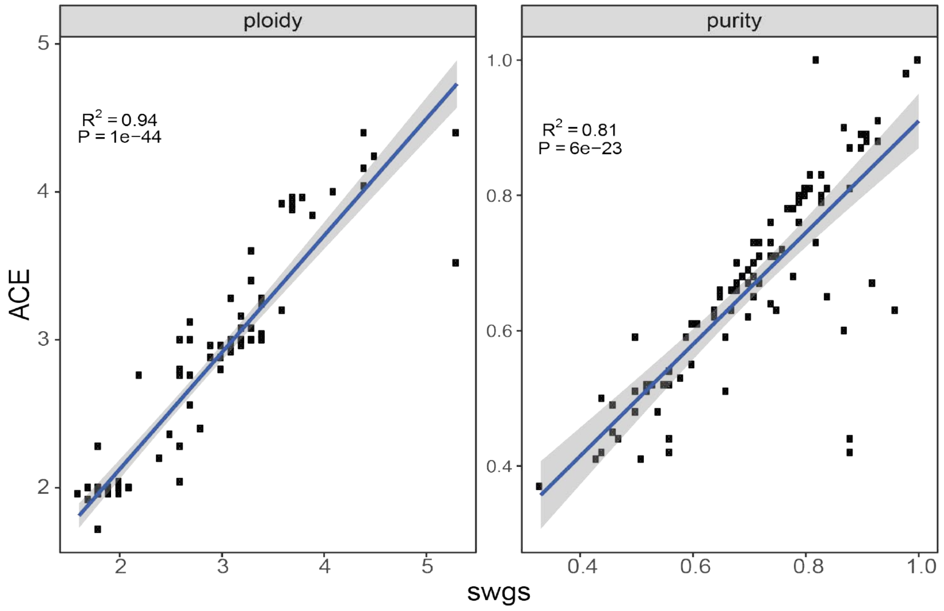

Figure S4

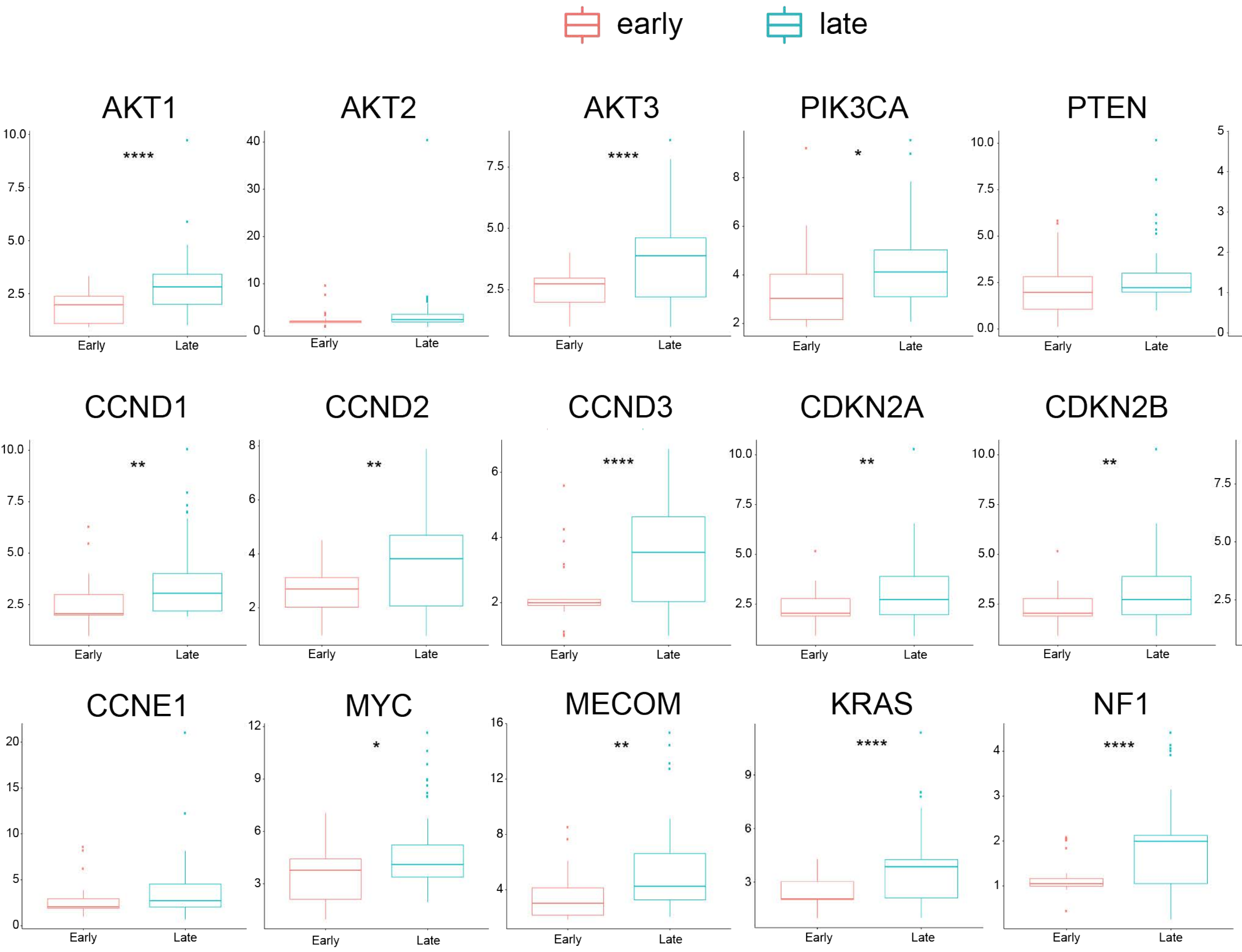


Figure S5

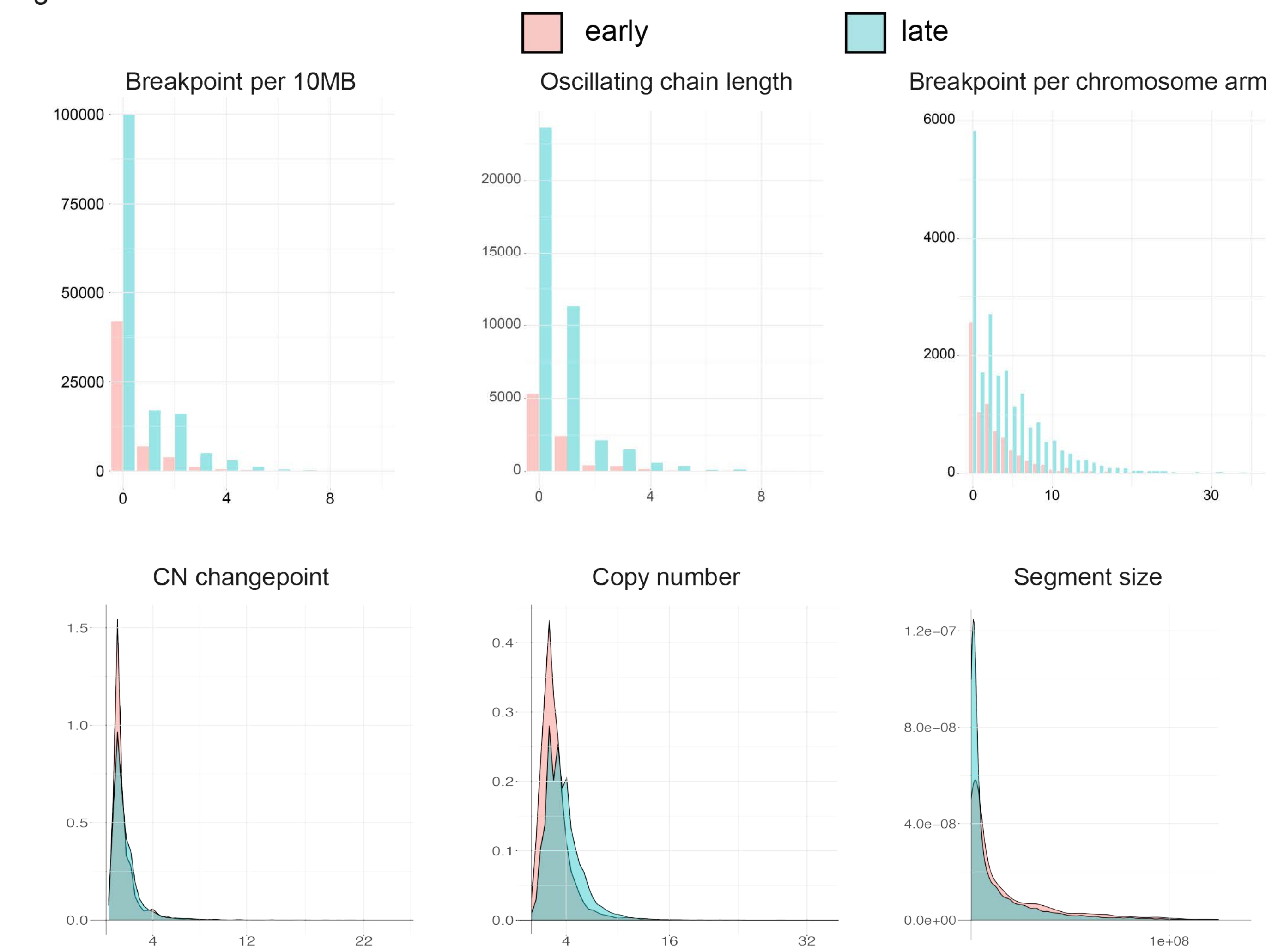
